# StACKER, A TOOL FOR SYSTEMS LEVEL ANALYSIS OF BASE STACKING IN NUCLEOTIDE-RICH STRUCTURES

**DOI:** 10.1101/2025.04.13.648419

**Authors:** Eric D. Sakkas, Daniel Krizanc, Kelly M. Thayer, Michael P. Weir

**Affiliations:** Department of Biology, Wesleyan University, Middletown, CT 06459, USA; Department of Mathematics and Computer Science, Wesleyan University, Middletown, CT 06459, USA; College of Integrative Sciences, Wesleyan University, Middletown, CT 06459, USA; Department of Chemistry, Wesleyan University, Middletown, CT 06459, USA

**Keywords:** Pi-stacking, ribosome CAR surface, protein translation, ribosome translocation, molecular dynamics, systems biology

## Abstract

Nucleic acid macromolecules can undergo significant structural changes—often mediated by nucleotide modification or molecular interactions. These alterations can be critical to their function. An ongoing challenge is to develop useful tools to quantify system-wide changes in nucleic acid-rich structures. We introduce StACKER, a robust Python package for observing conformational changes in a nucleic acid structure through its pi-stacking of base rings. StACKER creates System Stacking Fingerprints (SSFs) which highlight the landscape of pi-stacking throughout a molecule and can be used to show widespread conformational adjustments. Additionally, StACKER’s Pairwise Stacking Fingerprint (PSF) can further characterize pi-stacking in a select residue pair, showing how the effects of localized residue changes spread through a nucleic acid system. We apply StACKER to molecular dynamics (MD) simulations of a subsystem of the ribosome to reveal that alternative codons at the ribosome A-site and 3’ adjacent +1 codon position induce allosteric structure changes in the decoding center neighborhood, leading to a toggle between two conformational states. Through previous analysis of ribosome profiling data, we link these states to an observed fast/slow translation phenotype (Sun et al. 2024). We benchmark StACKER alongside other lenses for observing allostery to demonstrate its use in assessing allosteric shifts and its relevance to future structure-to-function studies.

## INTRODUCTION

Developing lenses to characterize fine-grain properties across molecular structures is important for systems-level understanding of structure conformations and behaviors. Nucleic acid structures are particularly sensitive to conformation changes because nucleotide structure is conducive to hydrogen bonding interactions, including Watson-Crick/non-Watson-Crick base pairing, and pi-stacking interactions (Bottaro et al. 2014). With appropriate fine-grain metrics, preferred conformational states can be identified via clustering approaches, and the effects across the system resulting from localized changes or mutations can be investigated to dissect allosteric mechanisms (Thayer et al. 2017). Preferred structures visited in the conformational space can be extracted from X-ray and cryo-EM data and conformational changes over time can be modeled using molecular dynamics (MD) simulations which now have increasingly accurate force fields (Maier et al. 2015). Utilizing MD to analyze the effects of residue changes is a “computational genetics” approach that allows us to build a mechanistic understanding of neighborhood behaviors (Barr et al. 2020; Scopino et al. 2020; Scopino et al. 2021; Dalgarno et al. 2022; Sun et al. 2024).

In nucleic acid-rich MD structures, a network representation of the system can be established using fine-grain metrics as edge values between residue nodes (amino acids or nucleotides) (Chen and Lai 2018; Dalgarno et al. 2022). The pairwise edge values can be defined by residue-to-residue geometric measurements (e.g. distances between each residue’s backbone atoms) or molecular interaction measurements (e.g. electrostatic, van der Waals, or hydrogen-bond). Here, we focus on another edge value: pairwise pi-stacking interactions (Martinez and Iverson 2012; Carter-Fenk and Herbert 2020). These are particularly prevalent in base stacking of nucleotide-rich structures (Bottaro et al. 2014).

Pi-stacking behavior is not directly captured by structure backbone properties (Scopino et al. 2020; Sun et al. 2024) or other pairwise interaction measures used for system-level analysis (Case et al. 2023). Incorporating system-level analysis of base stacking could reveal important behaviors. Pi-stacking, also known as π-π stacking or base stacking, can be leveraged to incorporate the nitrogenous bases into systems analysis. This electrostatic interaction occurs between two conjugated rings 3-4 Å apart (Bottaro et al. 2014). All nitrogenous bases of the ribosomal RNA contain conjugated rings, and base stacking is critical to a stable RNA/DNA structure (Sierański 2020). Conjugated rings have delocalized electron clouds above and below the ring with a partial negative charge, inducing a partial positive charge on the atoms of the ring. Pi-stacking can occur in multiple conformations, including face-to-face/parallel stacking (stable in hydrophobic environments), edge-to-face/T-shaped stacking (common in proteins), and parallel displaced stacking (common in nucleic acid structures). RNA chains exhibit parallel displaced or slightly tilted sandwich stacking (a hybrid between T-shaped and parallel stacking) between their nitrogenous bases—two conformations that induce a dipole across the ring and lead to a stabilizing, attractive interaction (Figure 1A, B) (Martinez and Iverson 2012; Carter-Fenk and Herbert 2020). Additionally, RNA chains can form pi-stacking interactions with pi-bonds on amino acid residues, such as those in phenylalanine, tyrosine, tryptophan, histidine, and arginine.

**Figure 1.**
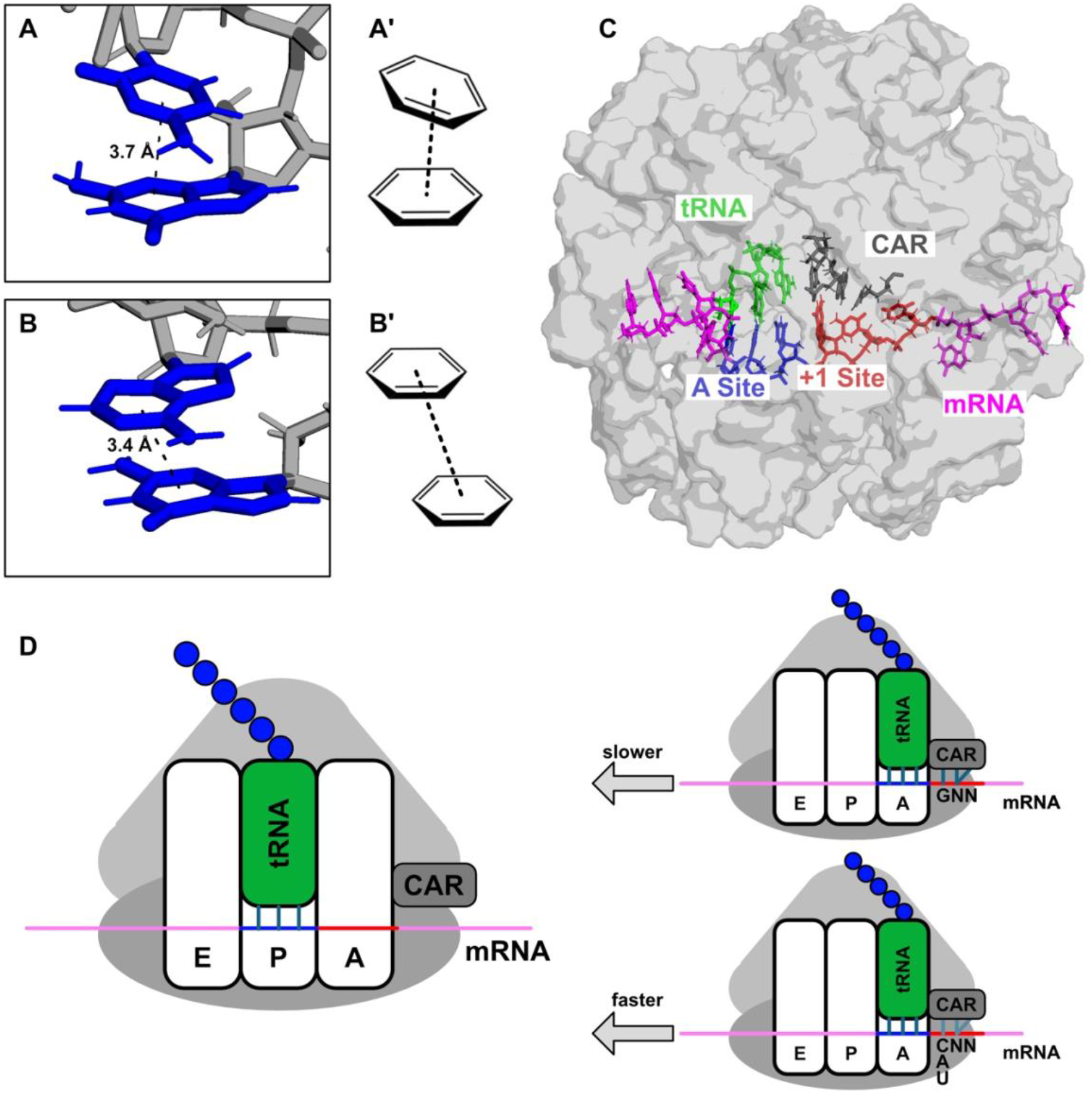
Pi-stacking of nucleotide base rings in the decoding center neighborhood. (**A**) Slightly tilted sandwich pi-stacking between a guanine and cytosine in the ribosome structure, visualized in PyMOL (Darden et al. 1993). The center of geometry (COG) distance is <4 Å, indicating pi-stacking. (**A’**) Schematic of tilted sandwich pi-stacking. (**B**) PyMOL visual of parallel displaced pi-stacking between guanine and adenine in the ribosome structure. (**B’**) Schematic of parallel displaced pi-stacking. (**C**) The subsystem of the yeast ribosome (494 residues in the decoding center) used to test StACKER. This decoding center neighborhood includes the mRNA, A-site codon, anticodon, +1 codon, and CAR. (**D)** Schematic of a +1GNN codon slowing mRNA threading, decreasing translation speed, as shown by ribosome profiling studies (Wu et al. 2019; Sun et al. 2024).

We have developed a software package StACKER (Stacking Analysis and Conformational Kinetics for Examining Residues) for systems and pairwise residue analysis of nucleic acid base stacking interactions. The package is Python-based and easily incorporates input trajectories from MD studies, allowing comparisons of molecular structure and dynamics using built-in K-means clustering and principal component analysis. We apply StACKER to an RNA-rich subsystem of the ribosome containing the A-site decoding center (Figure 1C) and show that changes in stacking interactions across the system can be used to demonstrate allosteric effects of localized mRNA nucleotide changes that are associated with reduced rates of ribosome translation.

## RESULTS AND DISCUSSION

### StACKER overview

StACKER is a Python software package with the following library dependencies: scikit-learn (Buitinck et al. 2013), NumPy (Harris et al. 2020), Pandas (Team 2024), matplotlib (Team 2024), seaborn (Team 2024), and mdtraj (Mcgibbon et al. 2015). The features are available through both Python and a command-line interface. The MD simulation input is processed using the Python package mdtraj, enabling support for multiple MD file formats, including DCD (CHARMM/NAMD), XTC/TRR (GROMACS), LAMMPS trajectories, and MDCRD files (AMBER) (Case et al. 2023). Alternatively, users can input PDB files, which combine both trajectory and topology information, and are outputted by most standard MD software. StACKER includes built-in data processing to fine-tune the MD input before analysis. These include trajectory filtering (e.g. by frame, residue, and atoms), distance calculation between residues, and filetype conversion.

We benchmarked StACKER on a subsystem of the ribosome containing the A-site codon, its tRNA anticodon, the 3’-adjacent +1 codon, and a sphere of surrounding residues Figure 1C). This subsystem also includes the ribosome’s CAR interaction surface which is hypothesized to tune translation rates through sequence-dependent H-bonding with the +1 codon (Sun et al. 2024) (Figure 1D). We performed MD simulations on four versions of the ribosome decoding center neighborhood, as outlined below, resulting in 3,200-frame trajectories for each construct (Figure 2A). Constructs differed by the codon identities in the mRNA A-site, tRNA anticodon, and +1 site, and were labelled accordingly. For example, a topology with a 5’-UAG-3’ tRNA, 5’-CUA-3’ A-site, and 5’-GCU-3’ +1 codon was denoted as tUAG_aCUA_+1GCU (Figure 2A; third topology from the left). Our previous analyses (Scopino et al. 2020; Scopino et al. 2021; Dalgarno et al. 2022; Sun et al. 2024) have shown that a +1 codon modification may impact residues many angstroms (Å) away, suggesting an allosteric effect. Finer-grain analysis of changes in stacking can reveal further insights into the allosteric mechanisms.

**Figure 2:**
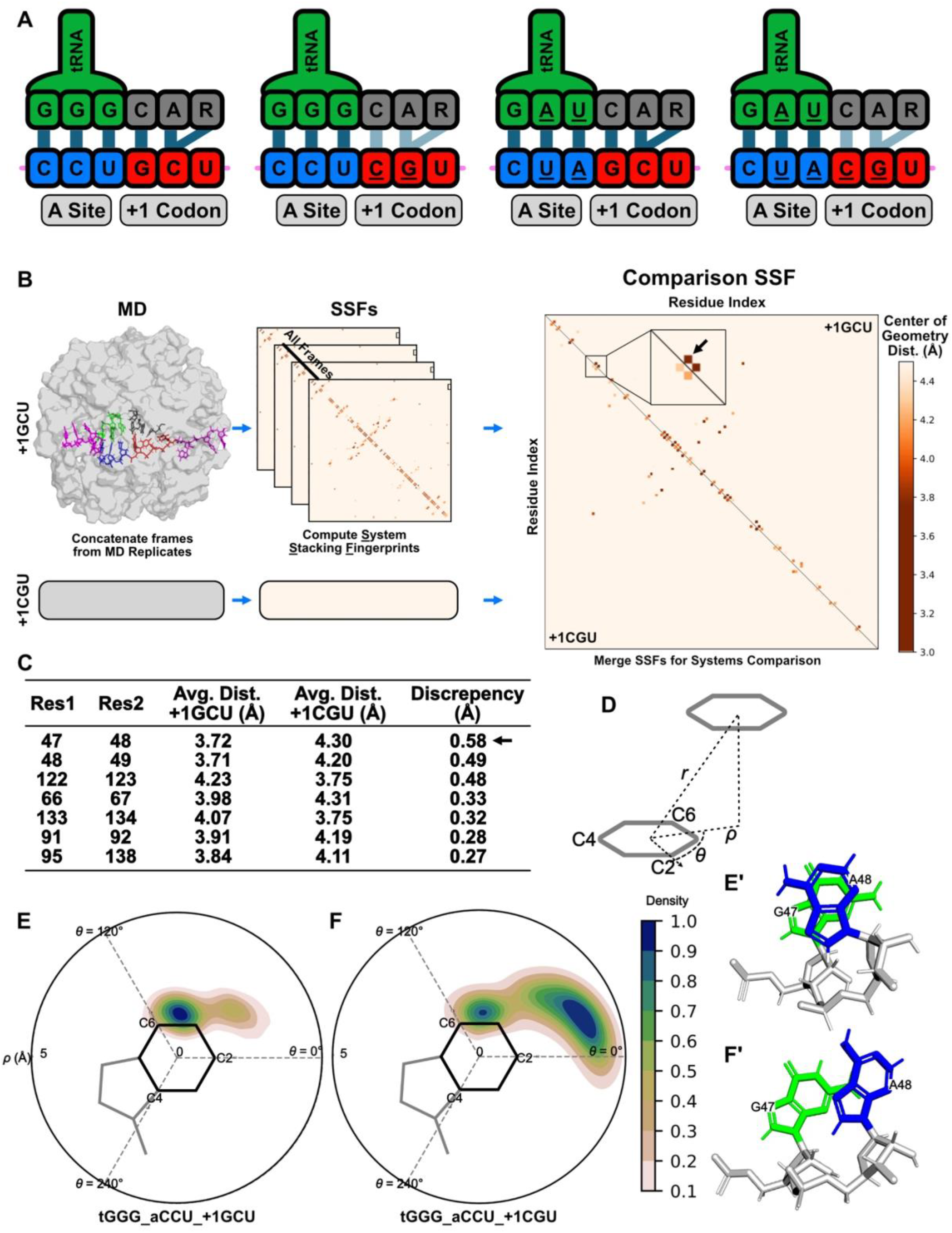
Stacking fingerprints of the CAR interaction surface. (**A**) The four MD structures used to benchmark StACKER. Nucleotide changes from the standard (left-hand) structure are underlined. From left to right: tGGG_aCCU_+1GCU, tGGG_aCCU_+1CGU, tUAG_aCUA_+1GCU, and tUAG_aCUA_+1CGU. (**B**) Pipeline for comparing tGGG_aCCU_+1GCU and tGGG_aCCU_+1CGU using System Stacking Fingerprints (SSFs). An SSF was made for each frame of the MD simulation and averaged into a single SSF for the whole trajectory. A comparison SSF shows the pi-stacking residue pairs in the +1GCU structure (top right triangle) and the +1CGU structure (bottom left triangle). The black arrow indicates residue pair 47/48, the pair with the largest change in pi-stacking. (**C**) Table showing the residue pairs whose pi-stacking properties (measured by avg. COG distance) changed the most between the tGGG_aCCU_+1GCU and tGGG_aCCU_+1CGU trajectories. (**D**) Measurements for quantifying pi-stacking. *r* denotes COG distance while *ρ* measures residue centeredness. (**E**) Pairwise Stacking Fingerprint (PSF) for Residues 47 and 48 in the tGGG_aCCU_+1GCU system shows a centered conformation, indicating pi-stacking of the two residues, as captured in PyMOL in **E’**. (**F**) PSF for Residues 47 and 48 in the tGGG_aCCU_+1CGU system shows that the residues adopt an often displaced conformation with low pi-stacking, as seen in **F’**.

### System stacking fingerprint (SSF)

System Stacking Fingerprints (SSFs) are matrix representations of the pi-stacking interactions throughout a molecule. As a proxy for pi-stacking, we used center of geometry (COG) distances between the base rings of nucleotide residues. We chose center of geometry rather than center of mass because a nucleotide’s COG is the center of its 6-membered ring and is therefore consistent between pyrimidine and purine residues. To create an SSF, we compute the COG distance *r*_*ij*_ for all pairwise combinations of residues *i* and *j* in a frame (Figure 2D). The *r*_*ij*_ value for a given pair of residues is averaged across all frames, resulting in an average COG distance for a given pair throughout the trajectory. A COG distance near 3.5 Å denotes pi-stacking, so residue pairs with an average *r*_*ij*_ value near 3.5 Å were colored within the output SSF. Alternatively, a user can select another COG distance threshold and color all residue pairs below this threshold.

To confirm COG distance *r* was an accurate proxy for stacking interactions, we examined residue pairs in our ribosome subsystem known to exhibit stacking: the tRNA codon/anticodon nucleotides. These stacked bases had an average distance between the base ring COGs of 4.39 ± 0.76 Å. The COG distances never fell below 3.27 Å suggesting that closer packing of the rings is energetically unfavorable.

As an additional confirmation that this was an accurate proxy metric for base stacking, 40 residue pairings with COG distance < 8 Å were randomly chosen from a frame of the tGGG_aCCU_+1GCU trajectory. The residue pairs were observed in PyMOL and manually labelled as either “Stacking” or “Not Stacking,” using criteria such as centeredness and parallel residue planes. To reduce bias, residue pairings were labelled in PyMOL without knowing their COG distance. Based on the results, a COG distance of < 4.5 Å was determined to indicate stacking, but the literature-accepted distance of 3.5 Å remains as a user option when creating SSFs.

We calculated SSFs for each of our four constructs using the 252 nucleic acid residues of the ribosome subsystem—restrained and unrestrained—while omitting any amino acid residues. Among all pi-stacking residue pairs (avg. COG distance < 4.5 Å) in the SSF for tGGG_aCCU_+1GCU (N = 106; Figure 2B top triangle, see below), most occurred near the diagonal of the SSF (N = 82, 77.4%), as these residue pairs were adjacent on the RNA chain, but many pi-stacking events occurred between nonconsecutive residues on the same chain or residues on different chains (N = 24, 22.6%).

### Comparisons of system stacking fingerprints

We compared SSFs between trajectories to observe the widespread pi-stacking changes throughout a system caused by the allosteric effect of a +1 codon change (+1GCU to +1CGU; Figure 2B). Since SSFs are symmetric across the SSF diagonal, comparison SSFs were made by filling the upper-right triangle of one matrix with the SSF from one trajectory, and the bottom-left triangle of the matrix with the SSF of another (Figure 2B). We complemented this with StACKER’s *compare* function, which outputs a rank-ordered table of the residue pairs with the largest change in average pi-stacking between trajectories (Figure 2C). These pairs were further explored with Pairwise Stacking Fingerprints (PSFs), as described below.

When comparing constructs tGGG_aCCU_+1GCU and tGGG_aCCU_+1CGU, which differed only at the +1 codon (the codon 3’-adjacent to the A-site codon), StACKER revealed many distant residue pairs whose pi-stacking quality significantly changed. Residues 47 and 48, a guanine and an adenine respectively, went from an average COG distance of 3.72±0.33 Å in the +1GCU structure to an average COG distance of 4.3±0.67 Å in the +1CGU structure, a significant (*t-test p <* 0.001) change of 0.58 Å (Figure 2C). This residue pair was 17.5 Å from the +1 codon (measured from the COG of the first nucleotide; Figure 3A), yet exhibited the largest change in pi-stacking. The pairs with the next largest stacking changes, residues 48/49 and residues 122/123, were 18.1 Å and 23.9 Å from the +1 codon respectively (Figure 3B, C). Pi-stacking changes far from the +1 codon site showed that the +1 codon replacement caused a structural change that permeated across the decoding center neighborhood.

**Figure 3:**
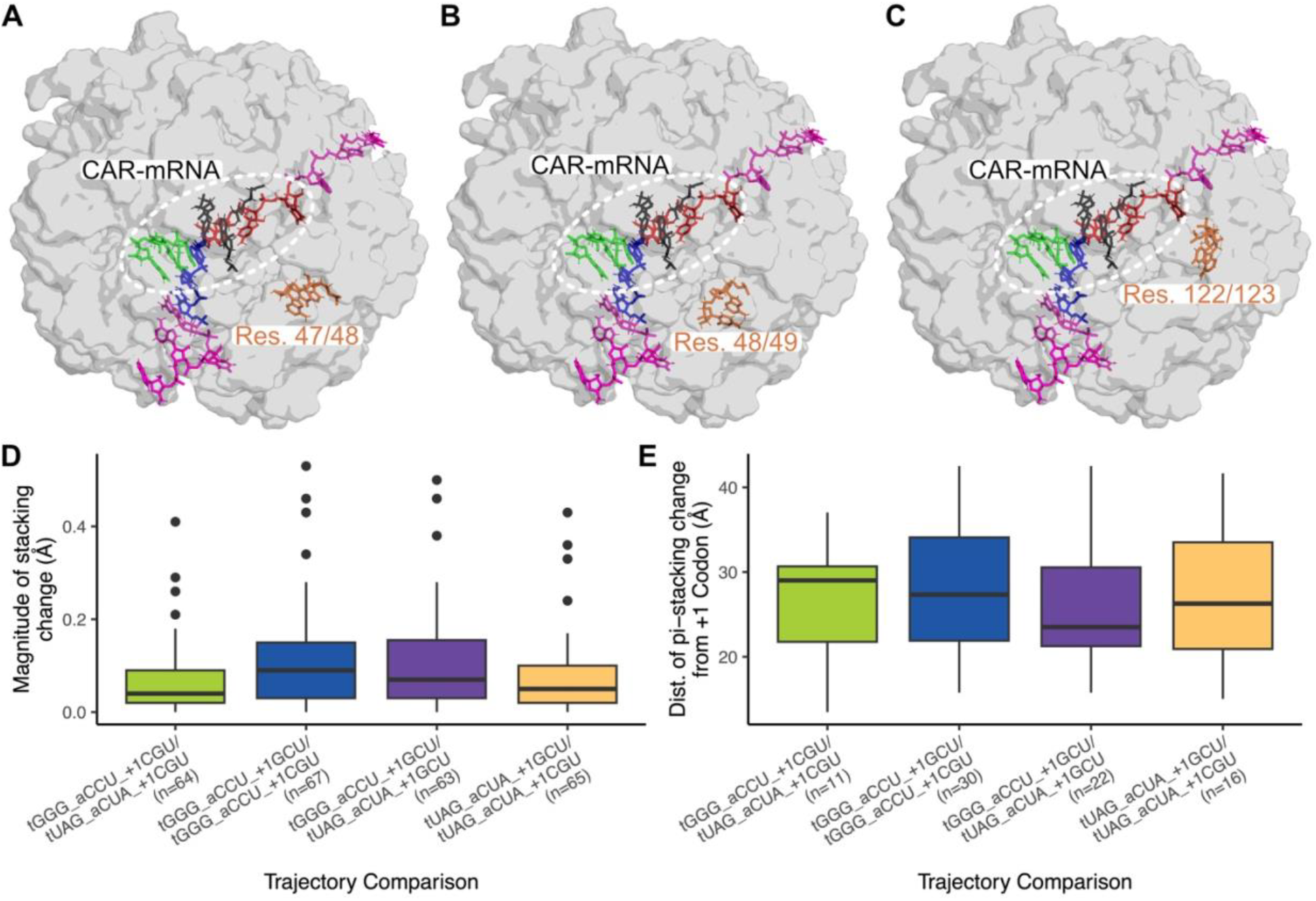
Pi-stacking changes permeate throughout the decoding center neighborhood. **(A-C)** The positions of the three residue pairs with the largest pi-stacking changes in the tGGG_aCCU_+1GCU/+1CGU structure (Figure 2C; Res. 47/48; 48/47; 122/123). The pi-stacking changes far from the +1GCU to +1CGU modification indicate the allosteric effect of the +1 codon. (**D**) Magnitudes of pi-stacking changes of residue pairs (with COG distance changes > Å) when comparing two trajectories. Included are all residue pairs with pi-stacking (COG distance < 4Å) in at least one structure of the comparison. (**E**) In all trajectory comparisons, pi-stacking changes occur throughout the structure, many far from the +1 codon modification. Plotted are distances between the COG of nucleotide 1 of the +1 codon and the closest residue of residue pairs with pi-stacking changes >0.1Å.

The allosteric impact of the +1 codon was maintained when we replaced the A-site codon identities in the tUAG_aCUA_+1GCU/ tUAG_aCUA_+1CGU comparison. Again, the structures differed only at the +1 codon, but the residue pairs with the largest pi-stacking changes were 194/197 (23.3 Å from +1 codon; data not shown), 122/123 (18.1 Å from +1 codon), and 290/291 (35.5 Å from +1 codon).

For several pairwise comparisons of structures (Figure 3D, E), we assessed the overall number of residue pairs whose pi-stacking COG distances changed by >0.1 Å. Across these comparisons, many of the implicated residue pairs had pronounced stacking distance differences (Figure 3D), and were located far from the +1 codon identity changes (Figure 3E), indicating allosteric effects throughout the structure. Consistent with our observations below, when trajectories were compared with the tGGG_aCCU_+1GCU trajectory, this revealed residue pairs with the largest COG distance changes, suggesting that the tGGG_aCCU_+1GCU structure had a distinct pi-stacking profile from the other trajectories.

### Pairwise stacking fingerprint (PSF)

Our analysis so far has focused on stacking behaviors from a system-wide perspective. We also built into StACKER the ability to examine the geometry of individual stacking events between the bases of pairs of residues. Using what we refer to as Pairwise Stacking Fingerprints (PSFs), we analyzed residue pairs with significant stacking changes between trajectories. Base stacking of residue pairs of interest was visualized using a polar coordinate representation, as in Bottaro et al. (Bottaro et al. 2014). For each frame, the COG of one residue in a residue pair was projected onto the plane of the other (Figure 2D), and the localization of one ring’s COG relative to the other was visualized as a heatmap. This helped differentiate between parallel displaced and sandwich pi-stacking, and these labels were confirmed by viewing the trajectories in PyMOL. A centralized COG, as in Figure 2E, denoted strong pi-stacking, while a delocalized position, as in Figure 2F, showed weaker stacking.

### Comparing pairwise stacking fingerprints of individual stacking events across systems

We created PSFs for the three residue pairs with the most pi-stacking change in our trajectory comparisons. The PSF for each residue pair was created using all frames (n=3,200) of that trajectory. The PSFs for the tGGG_aCCU_+1GCU/+1CGU comparison are shown in Figure 2E, F and Figure S1, while the PSFs for the tUAG_aCUA_+1GCU/+1CGU comparison are shown in Figure S2. Both comparisons revealed distinctive differences in stacking behavior in the different systems.

We observed the residue pairs with large COG distance changes between trajectories (discussed above) using StACKER’s Pairwise Stacking Fingerprints (PSFs). In the tGGG_aCCU_+1GCU and tGGG_aCCU_+1CGU comparison (a +1 codon modification) residues 47 and 48 exhibited the largest change in pi-stacking, an average COG distance change of 0.58 Å (Figure 2C). We characterized the pi-stacking change using the PSFs in Figure 2E, F. In the +1GCU structure, residues 47 and 48 have very little displacement, showing a strong pi-stacking conformation (Figure 2E’). In contrast, the +1CGU promotes a decoding center neighborhood structure that displaces residues 47 and 48, favoring a non-pi-stacked conformation for this residue pair (Figure 2F’).

Our PSFs also characterized the stacking changes in the residue pairs with the next largest pi-stacking change, residues 48/49 and 122/123 (Figure S1). Residues 48/49 (Figure S1A, B) displayed analogous characteristics to those in Figure 2E, F, with the +1GCU structure promoting pi-stacking and the +1CGU displacing the residues. Similarly, the two +1 codons led to different conformations in residues 122/123 (Figure S1C, D). In the +1GCU structure, residues 122 and 123 display two dominant conformations: one promoting pi-stacking and the other displaced, while the +1CGU structure promotes consistent pi-stacking between the two residues.

Exploring a different A-site codon, our tUAG_aCUA_+1GCU/tUAG_aCUA_+1CGU comparison showed that the allosteric impact of the +1 codon persists when we changed the A-site codon identities. While the magnitudes of discrepancies between COG distances were similar regardless of A-site sequences, the identities of the residue pairs with pi-stacking changes differed (Table S1). The residue pairs with the largest pi-stacking change in this comparison were residues 194/197, 122/123, and 290/291. The PSFs for these residues showed analogous results and are outlined in Figure S2.

### Systems comparison of individual fingerprints through K-means clustering and PCA

StACKER offers multiple methods for systems-level comparison of stacking fingerprints. We used StACKER’s built-in K-means algorithm to cluster system stacking fingerprints (SSFs) and identify structures with similar pi-stacking. For each construct, we created one SSF for each frame of a trajectory, totaling 3,200 SSFs per structure. Then, we gathered the SSFs from all four constructs into an input for K-means clustering, an input size of 12,800 SSFs to cluster, where the algorithm was blind to the source trajectory for each SSF.

We explored running the K-means clustering with different K values. Setting K=2 resulted in complete separation of the tGGG_aCCU_+1GCU trajectory from the three other trajectories (Figure 4A). This striking result indicated that the tGGG_aCCU_+1GCU construct exhibits a distinctive pi-stacking profile compared to the other structures, further supporting our hypothesis that changing the decoding center codon identities results in widespread conformational changes throughout the decoding center neighborhood. Increasing the number of clusters to K=8 partitioned the trajectories between clusters, though the tGGG_aCCU_+1GCU trajectory still separated independently into three clusters (Figure S3).

**Figure 4:**
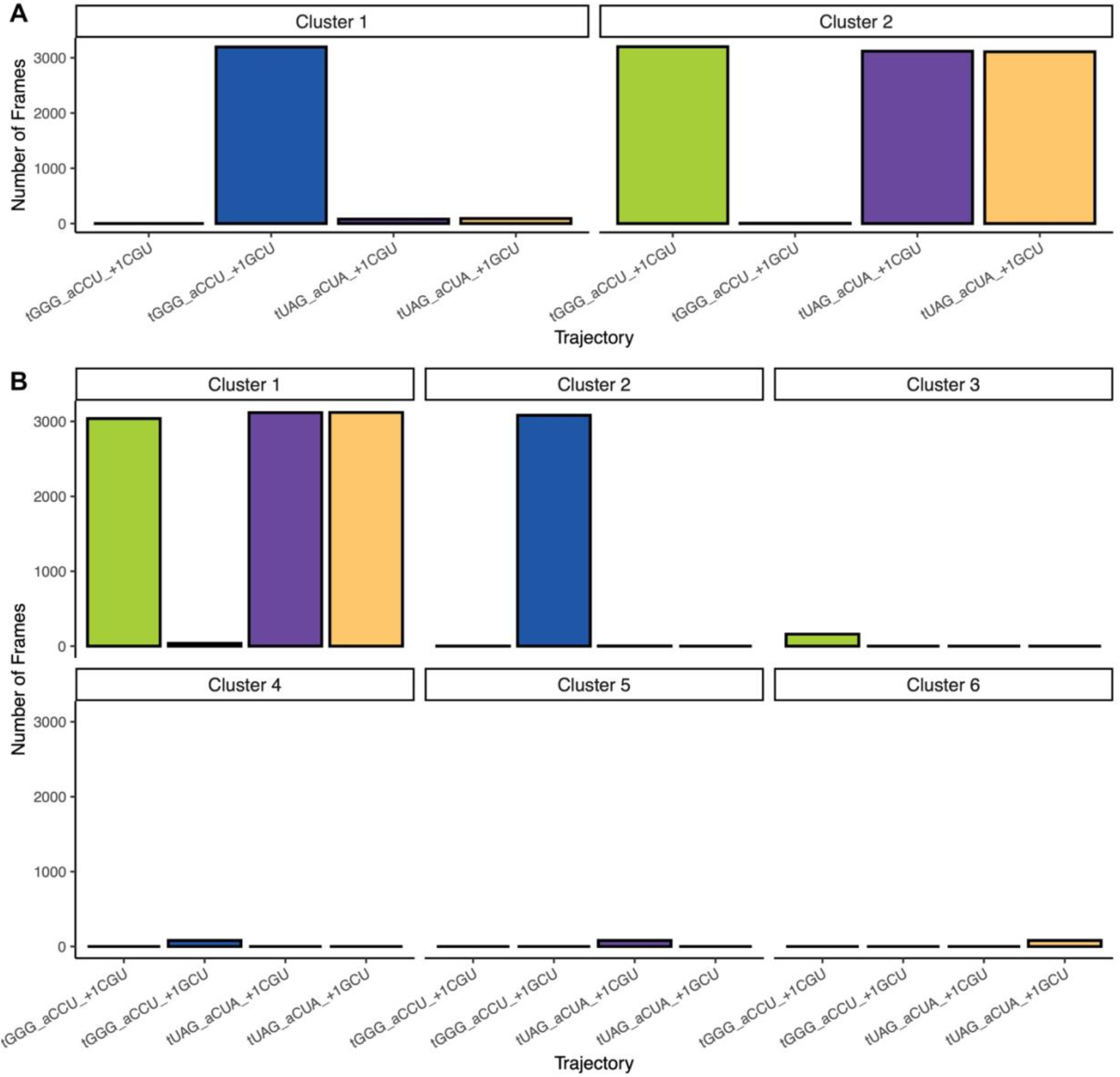
Systems comparison through K-means clustering. (**A**) K-means clustering (K=2) of all system stacking fingerprint (SSF) frames of all four decoding center constructs. The tGGG_aCCU_+1GCU structure clusters independently, suggesting system-wide differences in pi-stacking characteristics. (**B**) K-means clustering of MD frames using RMSD of the backbone atoms of the 321 unrestrained residues as the centroid distance measurement. At K=6, the tGGG_aCCU_+1GCU structure separates into an independent cluster, suggesting trajectory-specific preferred conformations.

Next, we used StACKER’s built-in principal component analysis (PCA) on the same frame-wise SSFs used in the K-means analysis. This allowed us to visualize the distinct pi-stacking characteristics of each frame on the Cartesian plane and observe variation in pi-stacking between trajectory frames. Principal Component 1 captured 9% of the data’s variability, while Principal Component 2 captured 7%, which provided enough separation to distinguish pi-stacking between systems. The PCA Plot in Figure 5A showed that frames originating from the tGGG_aCCU_+1GCU structure mainly occupied the space with Principal Component 1 > 0, while frames from the other three structures were mainly plotted with Principal Component 1 < This notable separation complemented our K-means results that the frames from tGGG_aCCU_+1GCU occupy their own cluster. Figure 5B colors the same PCA Plot by each frame’s K-means cluster identity from Figure 4A. We observed that the clusters generally occupy separate regions in the reduced dimensionality space, with Cluster 1 containing nearly all frames with Principal Component 1 > 0, which is associated with the tGGG_aCCU_+1GCU trajectory (Figure 5A). Thus, StACKER’s principal component analysis further supported our finding that the tGGG_aCCU_+1GCU has a distinct pi-stacking profile due to widespread structural differences compared to the other constructs.

**Figure 5:**
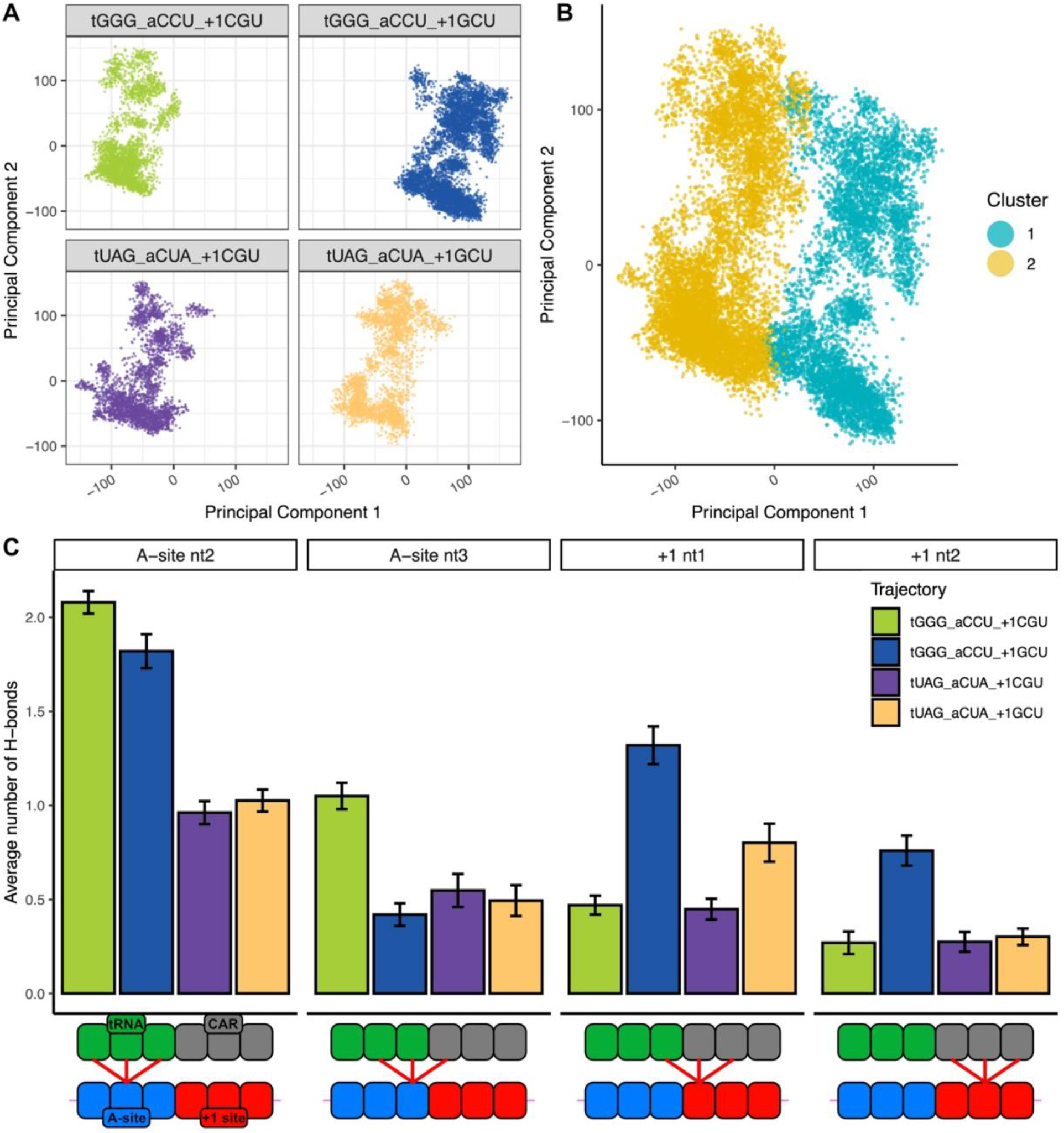
Independent methods support StACKER results. (**A**) PCA plot of SSFs facetted by trajectory. Each dot is an SSF of a single frame. The frames of the tGGG_aCCU_+1GCU structure mainly occupy the space where Principal Component 1 > 0, supporting a pi-stacking profile distinct from the other trajectories. (**B**) PCA plot with frames labelled by their K-means cluster in Figure 4A. The PCA plot coordinates accurately capture the same pi-stacking differences captured by K-means, as the clusters occupy distinct spaces in the plot. (**C**) Hydrogen bonding of A-site (nt2 and nt3) and +1 codon nucleotides (nt1 and nt2) with anticodon and CAR residues. The tGGG_aCCU_+1GCU structure displayed significantly stronger +1 codon/CAR hydrogen bonding.

### Validation of results with independent analyses

To confirm the unique structural characteristics of the tGGG_aCCU_+1GCU decoding center neighborhood, we applied multiple independent systems-level analyses. First, we complemented StACKER’s K-means analysis with an independent K-means analysis that clustered frames using RMSD (compared to reference centroids) of the Cartesian coordinates of backbone atoms of the 321 unrestrained residues. A cluster number of K=6 resulted in two majority clusters, one where the tGGG_aCCU_+1GCU separated into a distinct cluster, and another that contained the majority of frames from the remaining three trajectories (Figure 4B). As with our K-means clustering of system stacking fingerprints, this result supported our conclusion that the tGGG_aCCU_+1GCU structure creates a distinct decoding center neighborhood conformation. The complementary results show that StACKER adds to other system-wide measures a powerful method to characterize the nitrogenous base stacking at a systems level.

To further elucidate the mechanism by which the tGGG_aCCU_+1GCU structure differs from the other trajectories, we observed the average number of H-bonds formed between the tRNA/CAR and the mRNA. Consistent with our previous analyses, the tGGG_aCCU_+1GCU structure showed higher average number of H-bonds between the +1 codon and CAR than the other three structures (Figure 5C). The two +1CGU structures had less H-bonding with CAR than +1GCU. This difference in H-bonding was consistent with the separation of tGGG_aCCU_+1GCU in the K-means and PCA results. The identity of the A-site mRNA codon influences the H-bonding of the +1 codon with CAR (Sun et al. 2024). While the tUAG_aCUA_+1GCU structure had elevated +1 codon-CAR H-bonding compared to the +1CGU structures, the hydrogen bonding was still suppressed compared to tGGG_aCCU_+1GCU, and the structure clustered with the +1CGU trajectories in the K-means and PCA analysis. Therefore, our result that the tGGG_aCCU_+1GCU structure had higher H-bonding between the +1 codon and CAR than our other three trajectories is consistent with our previous analyses and, as discussed below, suggests a slower translation rate than the other tested A-site/+1 codon combinations.

Finally, we explored phenotypic differences between the four structures by using our previous analysis of ribosome profiling data (Sun et al. 2024). Ribosome profile densities provided a proxy for translation rate of the ribosome at particular A-site/+1 site codon identities. We found that NNU codons followed by +1GNN codons (where N represents any nucleotide identity) generally have much higher ribosome densities than when not followed by +1GNN. Our observations above that tGGG_aCCU_+1GCU trajectories are phenotypically distinct from tGGG_aCCU_+1CGU are consistent with these previous observations that aNNU_+1GNN codon pairs have higher ribosome densities and, therefore, slower translation. In contrast, our previous analysis of ribosome profiling data showed that compared to NNU, with NNA at the A site, the increases in ribosome densities for +1GNN (compared to when not followed by +1GNN) were much more muted, consistent with our observations above that tUAG_aCUA_+1GCU and tUAG_aCUA_+1CGU trajectories were phenotypically more similar in our testing with K-means clustering, PCA analysis, and H-bonding assessment.

## Conclusions

StACKER’s analysis of nucleic acid systems through their pi-stacking reveals subtle conformational changes and allows users to incorporate nitrogenous bases of nucleotides into systems comparisons. Our benchmarking of StACKER on the ribosome decoding center revealed an mRNA sequence-specific allosteric shift in the structure. We associated this shift with a translation speed phenotype consistent with our previous analyses (Scopino et al. 2020; Scopino et al. 2021; Dalgarno et al. 2022; Sun et al. 2024). Additionally, StACKER’s results aligned with multiple independent analyses, including hydrogen-bonding analysis, K-means clustering of backbone structures, and ribosome profiling. StACKER’s built-in downstream analyses, including K-means clustering and principal component analysis of system stacking fingerprints, and pairwise stacking fingerprints, provide a suite of robust methods for future systems comparisons of nucleic acid-rich structures (Joseph et al. 2021; Winkler et al. 2023; Dans 2024).

## MATERIALS AND METHODS

### The CAR interaction surface of the ribosome

We applied StACKER to an RNA-rich subsystem of the ribosome machine consisting of 494 residues in the ribosome’s A-site decoding center (Scopino et al. 2020; Scopino et al. 2021). This structure contained the A-site codon, A-site tRNA anticodon, a GCU +1 codon (the next codon to enter the A-site following translocation), and CAR (Figure 1C). CAR is an interaction surface of the ribosome comprised of yeast 18S rRNA C1274 and A1427—which we refer to by the corresponding 16S rRNA residues in *E. coli*, C1054 and A1196—and R146 in the ribosomal protein Rps3. The surface exhibits hydrogen bonding and pi-stacking with the +1 codon (Abeyrathne et al. 2016; Scopino et al. 2020; Scopino et al. 2021; Dalgarno et al. 2022). Our previous studies (Sun et al. 2024) of ribosome profiling data (Wu et al. 2019) have shown that the +1 codon-CAR interaction acts as a mechanism for tuning translation speed, with +1GNN codons slowing mRNA threading (Figure 1D). We have hypothesized that this phenotypic change is in part due to stronger hydrogen bonding between the +1GNN and CAR compared to other +1 codons (Scopino et al. 2020; Dalgarno et al. 2022; Sun et al. 2024).

Here, we expand this hypothesis: the +1 codon further tunes translation speed by inducing an allosteric structural change throughout the decoding center neighborhood. Changes in the pi-stacking landscape of the ribosome system can be used to demonstrate the allosteric effects of a localized single mRNA codon that have been associated with different conformations. The resultant structures have been further associated with a slow/fast translation phenotype (Sun et al. 2024).

### Molecular dynamics simulations

The MD simulations of the A-site decoding center were created and processed as outlined in our previous publications (Dalgarno et al. 2022). We used a 494-residue subsystem of the yeast ribosome undergoing translocation stage II (PDB ID: 5JUP) (Abeyrathne et al. 2016). This structure, shown in Figure 1C, includes a viral internal ribosome entry site (IRES) RNA that mimics the mRNA and tRNA. We included residues within a 40 Å sphere of C1054, the C of CAR, to capture surrounding residues that could be impacted by a +1 codon change. To minimize chain breaks, residues beyond this sphere were added, extending these chains to their natural ends. When the natural chain extended well beyond the sphere, it was instead capped with a chain terminal residue. 5’ phosphates capping chain ends were removed to align with the recommendations of AMBER and the forcefield used, ff14SB (Maier et al. 2015; Scopino et al. 2020). We applied a 20 kcal/mol·Å^2^ force to restrain the outer residues of the subsystem and simulate the sterics of the surrounding ribosome at translocation stage II.

Nucleotide identities in the A-site mRNA, A-site tRNA, and +1 mRNA codon were changed using AMBER22 tLEaP (Case et al. 2023) and 30 independent replicate trajectories (20 x 60 ns; 10 x 100 ns; sampled at 2 frames per ns) were performed for each version of the subsystem. We confirmed that the trajectories had settled by 20 ns using RMSD analysis and used the trajectory frames after 20 ns. Finally, we concatenated the replicates into a single, 3,200-frame trajectory for each construct, which were submitted to StACKER. Throughout this paper, we use “trajectory” to mean the 3200-frame concatenation of our 30 independent replicate trajectories. In total, four constructs were designed to test the versatility of StACKER, as shown in Figure 2A.

### K-means clustering and principal component analysis

K-means clustering into groups of similar trajectory frames was performed by concatenating per-frame SSF matrices from different trajectories and flattening each 2D matrix into a vector (e.g. 252 ×252 SSF was flattened into a 63,504-member vector). A frame’s SSF-vector was assigned to the closest centroid via a Euclidean distance measurement. This flattening of each frame’s pi-stacking data into a vector provides the basis for clustering.

We compared the SSF K-means clustering results to an independent K-means clustering analysis where distances to centroids were instead determined by RMSD using the backbone atoms of all unrestrained residues (n=321), as outlined in our previous analysis (Dalgarno et al. 2022). We performed both K-means analyses with 2-8 centroids (K=2, …, 8) and chose the optimal K using silhouette analysis (Rousseeuw 1987).

To further visualize the pi-stacking characteristics of each trajectory frame, we used principal component analysis (PCA) through the PCA class from the sklearn.decomposition module in Python. PCA is a linear dimensionality reduction that reduced the 63,504-dimension vectors to a 2-dimension space, allowing each frame’s SSF to be plotted on a Cartesian plane. We colored the PCA plot by both trajectory and K-means cluster to confirm that K-means analysis accurately captured the variability in pi-stacking characteristics between structures.

### Hydrogen bond quantification

The primary mechanism by which the +1 codon interacts with the CAR ribosome surface is through sequence-dependent hydrogen bonding with CAR, as described previously (Scopino et al. 2020). For each trajectory, we measured the H-bonding between CAR and the +1 codon as well as between the A-site codon and anticodon. Python scripts were used to parse the results of the AMBER cpptraj *hbond* function, which identifies in each trajectory frame H-bonds with an acceptor-to-donor heavy atom distance <3.0Å and an angle <135°.

## Code availability

StACKER is available under a GNU General Public License v3.0 on GitHub (https://github.com/esakkas24/stacker). Additionally, the software is thoroughly documented at https://esakkas24.github.io/. The GitHub and documentation pages contain several example files and pipelines to familiarize users with the software.

## ACKNOWLEDGEMENTS

We thank Pete Hwang, Luis Perez, Sonia Roberts, Erika Taylor, Kristen Scopino, Mitsu Raval, and Robert Lane for discussions, and Henk Meij for technical assistance with high-performance computing.

