## Supplemental Figures & Table for "StACKER, A TOOL FOR SYSTEMS LEVEL ANALYSIS OF BASE STACKING IN NUCLEOTIDE-RICH STRUCTURES"

### SUPPLEMENTARY DATA

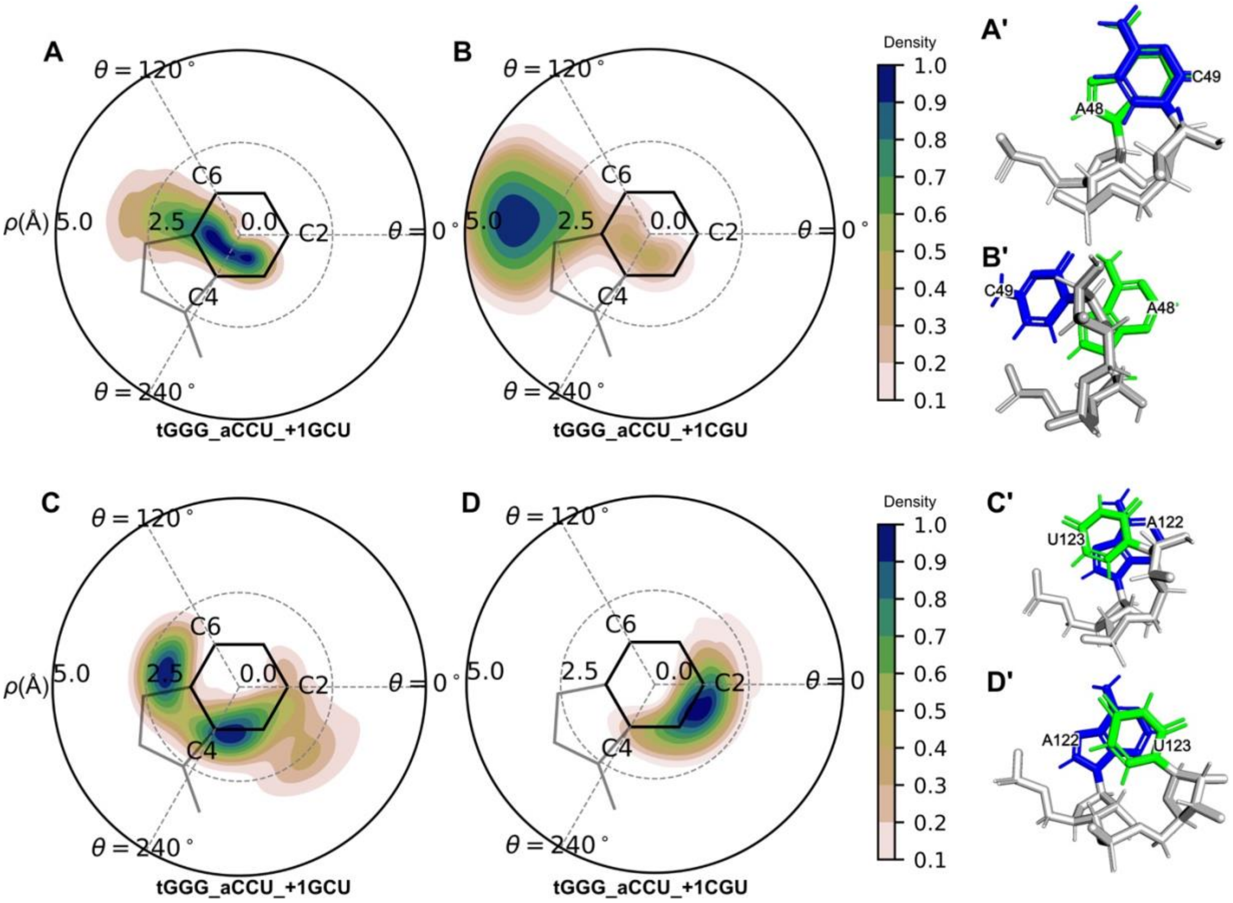


**Figure S1:** Pairwise stacking fingerprints (PSFs) showing pronounced changes in pi-stacking in comparison of tGGG_aCCU_+1GCU with tGGG_aCCU_+1CGU. (**A**) PSF of residue 48 and residue 49 in the tGGG_aCCU_+1GCU structure, showing a highly centralized conformation, indicating strong pi-stacking (captured in **A'**). (**B**) PSF of residue 48 and residue 49 in the tGGG_aCCU_+1CGU structure. The residue pair favors a displaced conformation, with low pi-stacking (see **B'**). (**C**) PSF of residue 122 and residue 123 in the tGGG_aCCU_+1GCU structure. The pair favors two conformations: one with parallel displaced pi-stacking, similar to the +1CGU structure, and another where the residue is either displaced with little to no pi-stacking or exhibiting pi-stacking with the 5-membered ring of the purine residue (see **C'**). (**D**) PSF of residue 122 and residue 123 in the tGGG_aCCU_+1CGU structure. The pair remains slightly displaced in a parallel displaced pi-stacking structure (see **D'**).

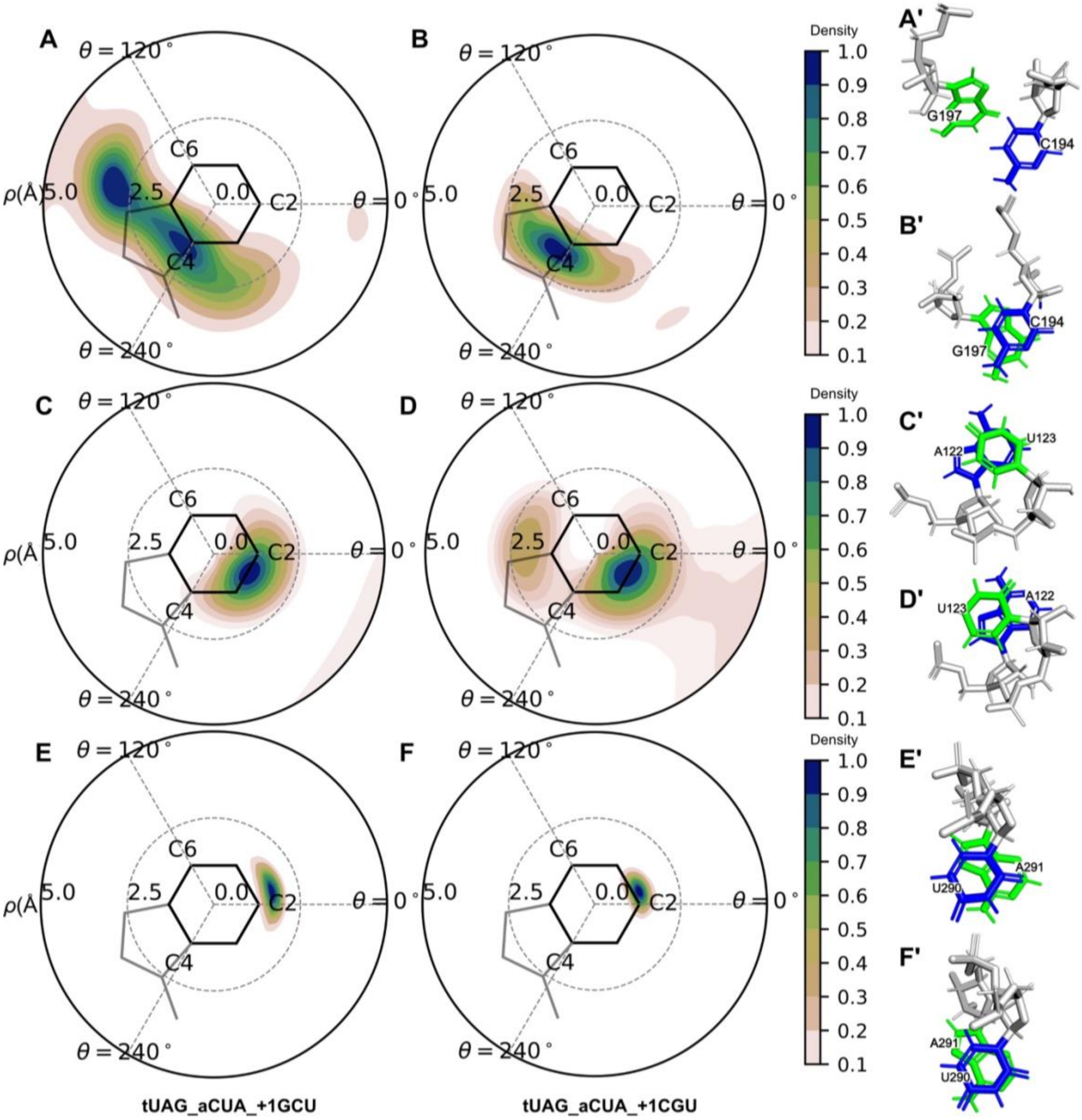


**Figure S2:** Pairwise stacking fingerprints (PSFs) of pi-stacking changes in comparison of tUAG_aCUA_+1GCU and tUAG_aCUA_+1CGU. (**A**) PSF of residue 194 and residue 197 in the tUAG_aCUA_+1GCU structure. The pair prefers two conformations, one pi-stacked and one decentralized. The latter is shown in (**A'**). (**B**) PSF of residue 194 and residue 197 in the tUAG_aCUA_+1CGU structure showing centralized pi-stacking (as illustrated in **B'**). (**C**) PSF of residue 122 and residue 123 in the tUAG_aCUA_+1GCU structure. The pair favors a parallel displaced pi-stacking conformation (see **C'**). (**D**) PSF of residue 122 and residue 123 in the tUAG_aCUA_+1CGU structure. The pair mostly maintains the pi-stacking of the +1GCU structure, but is occasionally displaced (as in **D'**). (**E, F**) The PSFs for Residues 290 and 291 in the tUAG_aCUA_+1GCU and tUAG_aCUA_+1CGU structures both show a focused strong pi-stacking conformation, with the +1GCU structure slightly more displaced. (**E'** compare with **F'**).

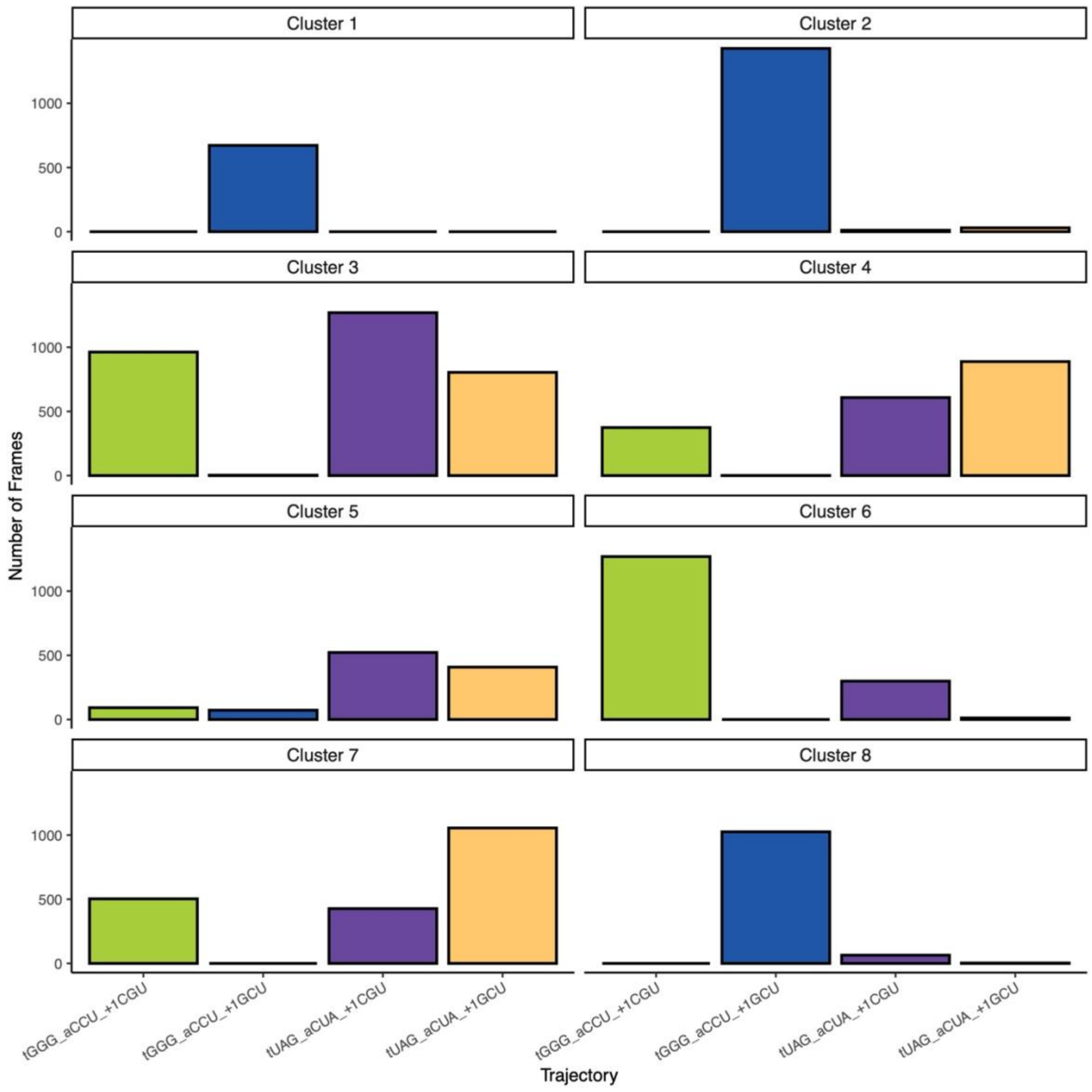


**Figure S3:** K-means analysis of SSFs with K=8. The tGGG_aCCU_+1GCU clustered independently, with 96.7% of clusters 1, 2, and 8 combined being comprised of frames from this trajectory. The remaining three trajectories were distributed among the remaining 5 clusters, with none of these clusters containing >90% frames from a single trajectory.

**Table S1:** Residue pairs by decreasing ΔCOG dist. in tUAG_aCUA_+1GCU/+1CGU comparison.

| Res1 | Res2 | Avg_Dist_tUAG_aCUA_+1GCU | Avg_Dist_tUAG_aCUA_+1CGU | Discrepancy |
|---|---|---|---|---|
| 194 | 197 | 4.41 | 3.98 | 0.43 |
| 122 | 123 | 3.8 | 4.16 | 0.36 |
| 290 | 291 | 4.2 | 3.87 | 0.33 |
| 72 | 73 | 4.05 | 3.81 | 0.24 |
| 95 | 138 | 4.11 | 3.94 | 0.17 |
| 79 | 80 | 4.11 | 3.98 | 0.13 |
| 97 | 136 | 3.89 | 3.76 | 0.13 |
| 57 | 58 | 3.86 | 3.73 | 0.13 |
| 104 | 106 | 4 | 3.87 | 0.13 |
| 138 | 139 | 3.85 | 3.73 | 0.12 |
| 59 | 60 | 3.85 | 3.97 | 0.12 |
| 81 | 82 | 3.9 | 4.02 | 0.12 |
| 180 | 181 | 4.09 | 3.97 | 0.12 |
| 61 | 62 | 3.78 | 3.89 | 0.11 |
| 170 | 172 | 3.96 | 3.85 | 0.11 |
| 101 | 102 | 3.58 | 3.69 | 0.11 |
| 183 | 184 | 4.03 | 3.93 | 0.1 |
| 201 | 202 | 3.89 | 3.98 | 0.09 |
| 5 | 6 | 3.62 | 3.71 | 0.09 |
| 163 | 164 | 3.81 | 3.72 | 0.09 |
| 58 | 59 | 3.91 | 3.99 | 0.08 |
| 103 | 104 | 3.99 | 3.91 | 0.08 |
| 76 | 78 | 3.8 | 3.87 | 0.07 |
| 85 | 86 | 3.68 | 3.61 | 0.07 |
| 108 | 109 | 3.73 | 3.66 | 0.07 |
| 172 | 173 | 3.77 | 3.7 | 0.07 |
| 177 | 178 | 3.43 | 3.5 | 0.07 |
| 11 | 12 | 3.83 | 3.77 | 0.06 |
| 161 | 162 | 3.72 | 3.66 | 0.06 |
| 141 | 142 | 3.92 | 3.98 | 0.06 |
| 160 | 161 | 3.89 | 3.95 | 0.06 |
| 195 | 197 | 3.95 | 4.01 | 0.06 |
| 156 | 157 | 3.56 | 3.51 | 0.05 |
| 97 | 98 | 3.99 | 3.94 | 0.05 |
| 43 | 44 | 3.5 | 3.55 | 0.05 |
| 133 | 134 | 3.78 | 3.73 | 0.05 |
| 54 | 55 | 3.44 | 3.4 | 0.04 |
| 112 | 113 | 3.75 | 3.71 | 0.04 |
| 136 | 137 | 3.76 | 3.72 | 0.04 |
